# Exploring the sensitivity limits of neuronal current imaging with MRI and MEG in the human brain

**DOI:** 10.64898/2026.02.17.706369

**Authors:** Milena Capiglioni, Davide Tabarelli, Stefano Tambalo, Federico Turco, Roland Wiest, Jorge Jovicich

## Abstract

**Introduction:** Conventional BOLD-fMRI relies on hemodynamic responses that are temporally and spatially indirect markers of neural activity. Developing alternative contrasts, sensitive to neuroelectrical phenomena, is a critical challenge in brain imaging. Spin-lock (SL) fMRI has shown promise in phantom studies for detecting magnetic field changes associated with neuronal activity, but its in-vivo sensitivity and practicality remain unclear. This study evaluated whether SL contrast can effectively detect and localize human neuronal activation, benchmarked against complementary functional modalities, magnetoencephalography (MEG) and 3T BOLD-fMRI, to assess the sensitivity of MR-based neuronal current imaging.

**Methods:** Thirteen healthy young volunteers underwent SL-based imaging during 8 Hz visual stimulation, along with BOLD and MEG acquisitions. Subjects viewed quadrant-checkerboard stimuli to elicit localized cortical responses. Two balanced SL contrast mechanisms, rotary excitation (REX) and stimulus-induced rotary saturation (SIRS), were employed. Postprocessing targeted stimulus-locked signal fluctuations using a regression-filtering-rectification strategy. Phantom experiments tested sensitivity and analysis pipeline performance.

**Results:** MEG revealed robust stimulus-locked responses in occipital cortex, with estimated local magnetic field amplitudes of ∼0.07 nT. Conventional BOLD-fMRI confirmed reliable hemodynamic activation. In contrast, neither balanced REX nor balanced SIRS produced consistent stimulus-related activation in vivo. Phantom experiments subsequently yielded detection thresholds of 0.2 nT for REX and 0.6 nT for SIRS, exceeding the MEG-estimated physiological field amplitudes.

**Conclusions:** Under the present experimental conditions, the tested spin-lock fMRI implementations did not achieve sufficient sensitivity for reliable in-vivo detection of neuronal magnetic fields at 3T. Phantom and MEG-based estimates indicate that physiological field amplitudes in the visual cortex lie below current detection limits. These findings establish quantitative constraints on direct neuronal current imaging with MRI and provide a benchmark for future methodological developments aimed at bridging electrophysiology and functional MRI.

**Key points:**

- We assessed spin-lock fMRI sensitivity using combined SL-fMRI, BOLD-fMRI, MEG, and phantom measurements during visual stimulation.
- MEG and BOLD-fMRI confirmed robust neuronal and hemodynamic activation in the visual cortex.
- SL-fMRI did not achieve reliable in-vivo detection of neuronal magnetic fields; phantom sensitivity limits exceeded MEG-estimated physiological field amplitudes.

## Introduction

The ability to non-invasively image neuronal activity with high spatial and temporal resolution remains a central challenge in neuroimaging. While blood oxygen level dependent (BOLD) functional MRI has revolutionized the study of human brain function, it gives only an indirect measure of neural activity through neurovascular coupling [Logothetis, 2008]. In contrast, electroencephalography (EEG) and magnetoencephalography (MEG) measure neuronal currents directly via electric and magnetic fields, but are limited by poor spatial resolution and source ambiguity due to the ill-posed inverse problem [Ahlfors et al., 2010; Cicmil et al., 2014; Gross et al., 2013; Hämäläinen et al., 1993; Michel and Murray, 2012].

To address this gap, researchers have explored non-BOLD MRI methods aimed at directly detecting neuronal magnetic fields. Early efforts focused on capturing small neuronal field-induced phase shifts in gradient-echo signals [Bodurka and Bandettini, 2002; Petridou et al., 2006], though in-vivo validation proved difficult [Chu et al., 2004; Parkes et al., 2007]. More recently, a 2D line-scanning method claimed high-temporal-resolution detection of neuronal currents, but initial reports were later retracted, and findings could not be reproduced [Choi et al., 2024; Phi Van et al., 2024].

Another emerging alternative is spin-lock functional MRI (SL-fMRI). First proposed in 2008[Witzel et al., 2008], SL-fMRI exploits the resonance interaction between spin-locked magnetization and oscillating neuronal magnetic fields to produce functional contrast. Because the signal depends on the initial phase of the neuronal field at the onset of the spin-lock pulse[Capiglioni et al., 2022], on-resonance SL-fMRI acquisitions are expected to exhibit increased signal variance compared with off-resonance acquisitions, allowing separation from conventional BOLD effects. Phantom studies have demonstrated that SL-fMRI can detect neuron-like magnetic field fluctuations with sub-nanotesla sensitivities[Halpern-Manners et al., 2010; Sveinsson et al., 2020]. However, in vivo application of SL-fMRI remains limited. Promising results have been reported in epilepsy, where neuronal activity is expected to be stronger[Kiefer et al., 2016; Unger et al., 2021], while studies in healthy volunteers have shown only group-level evidence during visual stimulation[Capiglioni et al., 2025; Truong et al., 2019], with limited or absent detection at the individual level[Capiglioni et al., 2025]. These findings highlight persistent questions regarding in-vivo sensitivity, robustness to field imperfections, and the ability of SL-fMRI to resolve physiologically relevant neuronal signals.

Two implementations of SL-fMRI have emerged: Stimulus-Induced Rotary Saturation (SIRS)[Witzel et al., 2008] and Rotary Excitation (REX) [Gram et al., 2022; Truong et al., 2019]. SIRS encodes signal changes via longitudinal magnetization after a tip-up pulse, whereas REX skips the tip-up pulse, and reads out after a slice-selective excitation applied right after the long SL pulse, offering potentially higher sensitivity but lower signal amplitude and greater susceptibility to field imperfections. Composite and balanced SL preparations, adapted from T1ρ mapping protocols [Wáng et al., 2015; Witschey et al., 2007], have recently improved robustness against B0 and B1 inhomogeneities [Capiglioni et al., 2022; Gram et al., 2022], though their performance in detecting in-vivo neuronal signals has yet to be established.

In this study, we evaluate and compare the functional sensitivity of both balanced SIRS and REX SL-fMRI sequences during visual stimulation in healthy volunteers using a 3T clinical scanner. We used BOLD-fMRI and MEG as benchmark modalities, while phantom experiments with identical acquisition protocols provide in-vitro sensitivity estimates, helping to contextualize in-vivo findings.

## Materials & Methods

The study was designed as a three-stage multimodal validation framework. First, in-vivo SL-fMRI, BOLD-fMRI, and MEG data were acquired during visual stimulation to assess functional sensitivity under realistic experimental conditions. Second, MEG recordings were used to estimate stimulus-locked neuronal magnetic field amplitudes in the visual cortex. Third, dedicated phantom experiments employing identical acquisition and analysis pipelines were conducted to quantify the minimum detectable field amplitudes for each spin-lock implementation. This integrated approach enabled direct comparison between physiological signal magnitudes and sequence-specific sensitivity limits.

### Human experiments

Thirteen healthy participants (mean age 26.1 ± 5.9, range 20-43 years) underwent scanning on a 3T MRI scanner and magnetoencephalography. The study was approved by the Ethical Committee of the University of Trento, and all volunteers provided written informed consent prior to participation in the study. All participants had normal, or corrected to normal, visual acuity. The visual stimulation protocol was programmed in Psychtoolbox-3 (http://psychtoolbox.org/) and eye-tracker measurements were acquired in both modalities during the experiment. The MEG was performed at least three days before MRI to avoid the influence of residual magnetization [Gross et al., 2013]. For both modalities, the participants viewed a checkerboard quadrant stimulus flickering at 8Hz alternating with a rest period with black screen. The next subsections describe the details of the experiments and analysis for both modalities.

### MEG Data Acquisition

We recorded MEG data in a magnetically shielded room using a 306-channel Neuromag VectorView system (204 planar gradiometers, 102 magnetometers; MEGIN, Helsinki, Finland). Signals were sampled at 1 kHz (low pass filter at 0.3 Hz and high pass filter at 330 Hz). Participants viewed flickering checkerboards (8 Hz) while maintaining central fixation during alternating 8 s stimulation and 8 s rest periods. The first 18 flickering periods occurred in the lower left and the second 18 in the lower right visual field (Figure 1a). At each transition between stimulation conditions, a 2 s interval allowed participants to blink. For each subject, we repeated the full experiment twice. During data acquisition, we continuously monitored head position using five head position indicator (HPI) coils. Before preprocessing, we realigned the head position to a reference common position using the Signal Source Separation method (SSS)[Taulu and Kajola, 2005], minimizing potential errors due to head movements. We recorded eye movements monocularly at 1 kHz using an MEG compatible eye tracking system to subsequently reject eye-movement related artifact.

**Figure 1:**
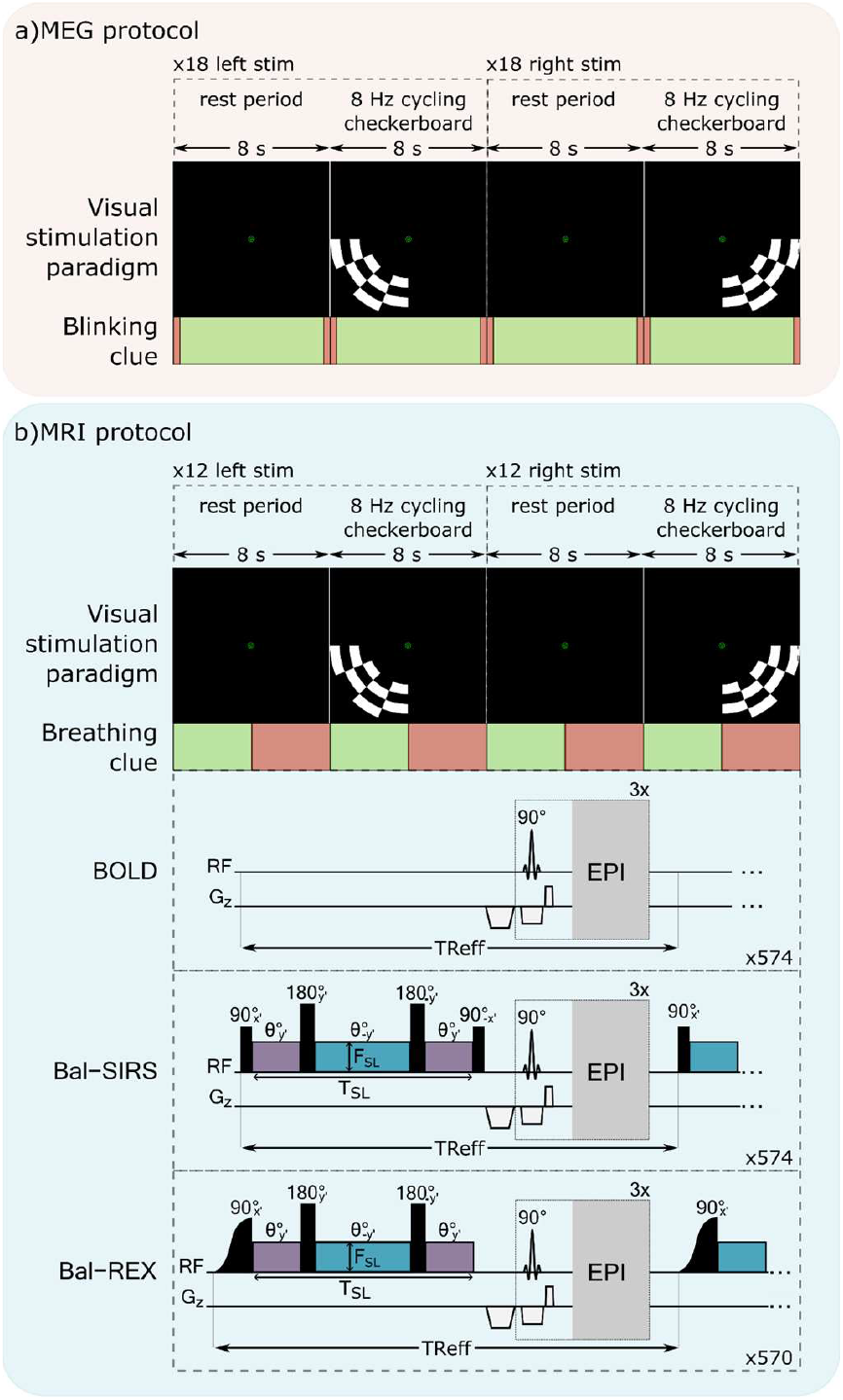
MEG and fMRI acquisition and visual stimulation protocols. a) Experimental paradigm for MEG. 8-second blocks of 8Hz checkerboard quadrant stimuli and 2-second blinking period indicated by fixation point. b) Experimental paradigm for functional MRI (three axial AC-PC slices covering the primary visual cortex). 8-second blocks of 8Hz checkerboard stimuli. Breathing periods of 4 s indicated by color change of the fixation point. MRI acquisition includes BOLD (equal timing to SIRS, omits SL preparation), Balanced SIRS, and Balanced REX. θ represents the induced rotation angle in each SL segment. A crusher gradient along z is applied after SL, followed by 3 consecutive EPI readouts.

### MEG Preprocessing and Analysis

We preprocessed and analyzed MEG data using MNE-Python[Gramfort et al., 2014]. This involved identification and removal of noisy channels and artifact-contaminated epochs, rejecting and correcting residual ocular and muscle artifacts with independent component analysis, and computing power spectra using a Discrete Prolate Spherical Sequences based multitaper method (4–42 Hz; epochs length: 4 sec; bandwidth: 2 Hz). Power spectra were computed and analysed, for each subject, at sensor level, as overall power spectral density and a time frequency map, with a 20 ms resolution.

We estimated the source of activation in the brain using a Dynamic Imaging of Coherent Sources approach[Gross et al., 2001]. The Fourier parameters were the same, leading to a spectral resolution of 0.25 Hz. We based cortical reconstruction on individual anatomical models derived from each participant’s T1-weighted MRI and FreeSurfer surfaces. We then computed the ratio of activity between stimulation and rest, and tested it statistically using a non-parametric permutation statistics approach to provide significant activation maps at the source level. The group source level activation maps were obtained from 11 out of 13 participants. One subject was excluded due to unsuccessful anatomical reconstruction from the T1-weighted scan, and another due to incorrect stimulation timing. The latter subject has been excluded also from the power spectra analysis

Finally, we estimated local magnetic field amplitudes in the occipital cortex from MEG recordings following the approximate approach reported by [Bodurka and Bandettini, 2002]. Starting from the magnetic field measured on a selected set of lateralized occipital sensors, each participant’s MEG data was band-pass filtered around the frequency of interest *f* using a narrow bandwidth (Δf = 1 Hz). We then computed the Hilbert envelope at the selected frequency and estimated the local magnetic field amplitude by multiplying the average Hilbert envelope by a factor of 10^3^. This provides an approximate conversion from sensor-level measurements to cortical field strength. We applied this procedure separately for left- and right-field stimulation conditions and subsequently averaged across subjects to obtain group-level estimates.

Further details regarding acquisition, preprocessing, and modelling procedures appear in Supporting Information 1.

### MRI Data acquisitions

#### In-vivo MRI experiments

We acquired MRI data on a 3T clinical scanner (MAGNETOM Prisma, Siemens Healthineers AG, Erlangen, Germany) equipped with a 64 channel head-neck receive coil. After standard T1-weighted MPRAGE images for anatomical reference, functional MRI data was acquired while participants performed a passive visual task designed to match the MEG experiment, with timing adjusted to fit MRI acquisition constraints. The visual stimuli were identical to those used in MEG: a flickering black-and-white checkerboard presented in either the lower left or right visual field. For MRI, each checkerboard was displayed for 8 s at 8 Hz, alternating with 8-s blank screens. The first 12 stimulation blocks targeted the lower left visual field, followed by 12 blocks in the lower right. A central fixation cross was displayed throughout the experiment, changing color every 4 s to cue breathing (Figure 1b). The controlled breathing was to ensure that the breathing frequency didn’t match the stimulation frequency (1/16 = 0.0652 Hz), and that it could therefore be eliminated by high pass filtering.

This paradigm was repeated for three functional MR sequences: balanced SIRS (Bal-SIRS)[Capiglioni et al., 2022; Witzel et al., 2008], balanced REX (Bal-REX) [Gram et al., 2022; Truong et al., 2019], and conventional BOLD-fMRI (Figure 1b). All sequences used a shared EPI readout covering the primary visual cortex with three AC-PC oriented axial slices (matrix size = 64 × 64 × 3; voxel size = 3.4 × 3.4 × 5 mm^3^; flip angle = 90°; TE = 29.8 ms; bandwidth = 1950 Hz/pixel, slice gap = 5 mm). For SIRS and REX, the SL frequency was set at 16 Hz for on-resonance condition and at 36 Hz to have an off-resonance control condition expected to give no functional sensitivity. Bloch simulations of the SL effect [Capiglioni et al., 2022; Capiglioni et al., 2023] were used to determine the optimal spin-lock duration (TSL), which was fixed at 112 ms for all acquisitions and frequencies, as this value maximized signal variation while maintaining consistent T_1_ρ weighting across conditions (see supporting information 2). The BOLD fMRI sequence used the same repetition time (TR) of 334 ms as the SL acquisitions, leaving the spin-lock preparation period empty to ensure comparable timing and signal evolution between sequences. Each acquisition began after 12 dummy scans, yielding a total duration of 6.4 min per sequence.

#### Phantom MRI experiments

To estimate the sensitivity limit for the Bal-SIRS and Bal-REX sequences, we used an electrical phantom previously described by [Capiglioni et al., 2022]. The phantom consisted of a 17-cm acrylic sphere filled with a 0.07 mM MnCl_2_ saline solution. A loop coil inside the phantom is connected to a function generator and generates oscillatory magnetic fields oriented parallel to the main static field *B*_O_. Figure 2a shows a schematic of the phantom setup.

**Figure 2.**
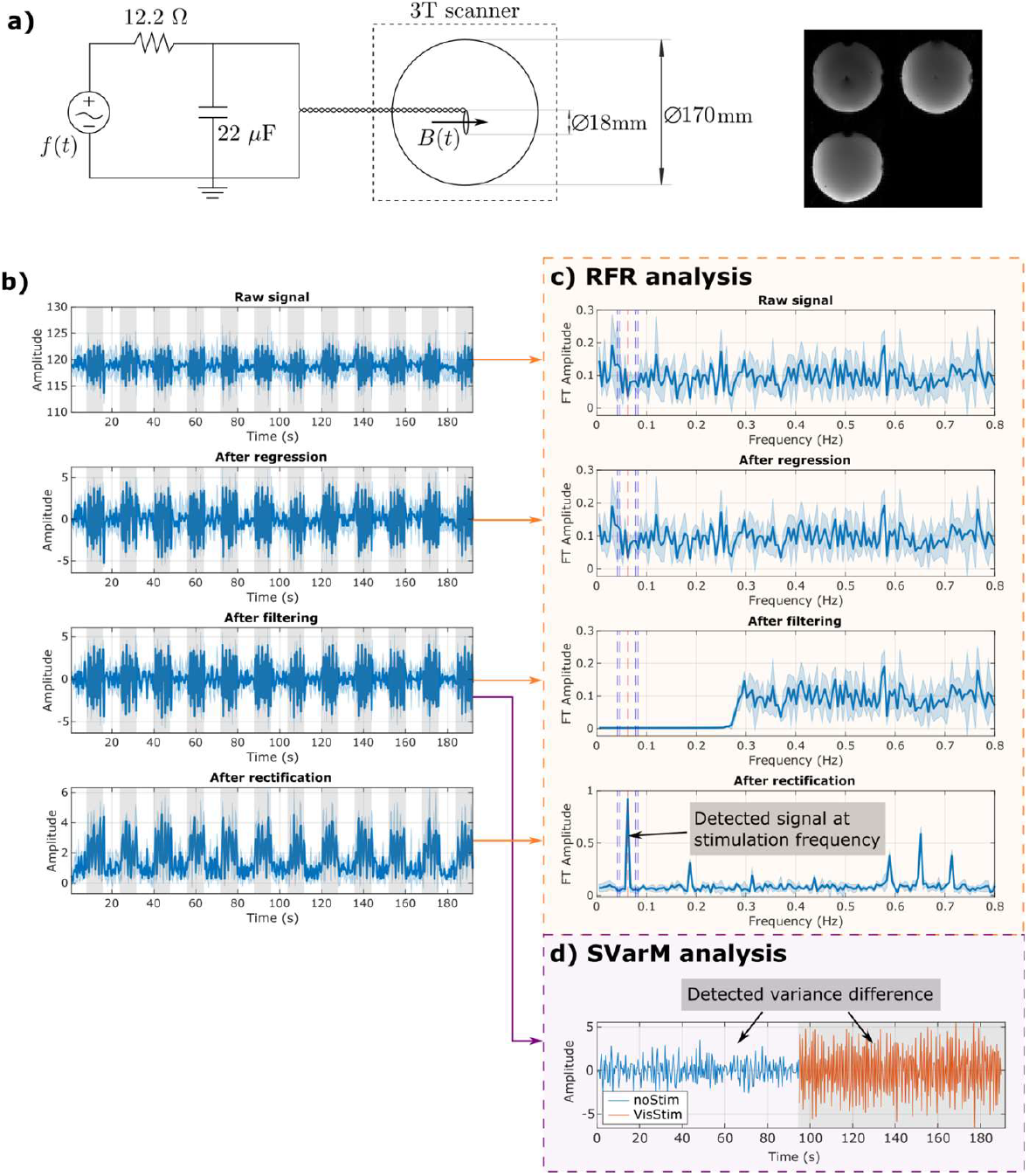
Postprocessing pipeline validation on phantom data. a) Diagram of the electrical phantom setup. The field inside the loop B(t) is calculated through the Biot Savart Law. Bal-SIRS image of the phantom is displayed on the right. b) Signal time course for the SL acquisition and output after regression, filtering, and rectification steps. c) Outputs of the RFR procedure in the frequency domain: the rectified signal highlights the detection at the block-stimulation frequency of 0.0625 Hz. d) SVarM tests variance differences of the on and off stimulation periods.

The input waveform (Figure 2b) reproduced the expected neuronal magnetic field during the visual stimulation paradigm used in the human experiment. During each 8 s stimulation period, we applied a sinusoidal current described by *A sin* (*ϕ* + *ωt*), where the initial phase *ϕ* was fixed at 0°, the angular frequency *ω* at 16 Hz, and *A* denotes the peak-to-peak voltage amplitude in mVpp. We performed two phantom experiments. First, to demonstrate the feasibility of detecting spin-lock fMRI contrast and to validate the analysis pipeline, we applied a high-amplitude input signal (A = 50 mVpp) to generate a strong oscillatory magnetic field. Second, to quantify the sensitivity limit, we systematically varied the input amplitude between 2 and 50 mVpp. The corresponding magnetic field amplitude, estimated using the Biot–Savart law, ranged from approximately 2.8 nT to 9 nT. For both experiments, because the repetition time (TR) was not an integer multiple of the expected evoked neuronal response (1/16 s), successive measurements sampled a slightly different initial phase of the oscillation.

#### MRI data analysis

Image preprocessing for in-vivo acquisitions included motion correction using MCFLIRT (FSL 6.04) [Jenkinson et al., 2002] and within-subject coregistration with the high-resolution image using SPM. For both the phantom and human data, the SL post-processing combined elements of both the Regression-Filtering-Rectification (RFR) method [Truong et al., 2019] and Statistical Variance Mapping (SVarM)[Coletti et al., 2021], with adaptations to accommodate the present acquisition protocol. The RFR procedure first cleans the data from low-frequency drifts and physiological nuances (mainly BOLD) and highlights the extra variance expected during stimulation. In the regression step, each voxel time course was modeled using the mean signal, a slow drift term, and for the in-vivo experiments, a regressor corresponding to the hemodynamic response function at the stimulation frequency calculated as 1/16 s = 0.0625 s. A high-pass filter was then applied to remove low-frequency and residual hemodynamic components (Figure 2b). The filter cutoff was defined at 0.02 Hz, right above the controlled breathing frequency. The RFR procedure rectified voxel time courses, converting random-phase signals into cyclic signals at the stimulation frequency. Contrast maps were computed as the power of the rectified signal at 0.0625 Hz (the block frequency). While RFR effectively isolates the periodic components associated with the task paradigm, it does not, by itself, provide voxel-wise statistical inference. To address this, SVarM was subsequently applied to an intermediate RFR output (specifically, the regression-filtered (RF) data) to perform statistical variance comparison with a Levene’s test between stimulation and baseline blocks (Figure 2d).

In the phantom this full analysis pipeline was applied to Bal-Rex and Bal-SIRS acquisitions measured as a function of the target field amplitude to determine the sensitivity limit. In vivo, it was applied to five acquisitions: on-resonance Bal-REX (16 Hz), off-resonance Bal-REX (36 Hz), on-resonance Bal-SIRS (16 Hz), off-resonance Bal-SIRS (36 Hz), and conventional BOLD EPI.

To assess the robustness of the analysis with respect to parameter selection, we repeated the complete processing pipeline under multiple configurations. Specifically, we evaluated alternative high-pass filter cutoffs (up to 0.3 Hz), applied image realignment using both SPM and FSL, and tested different temporal selections for the RFR analysis, including restricting stimulation and baseline blocks to their central 6 s, their initial 4 s, or the full block duration. We additionally varied regression parameters, including regressing or not the hemodynamic response function and modeling or omitting slow signal drifts. Unless otherwise specified, results are reported using the primary analysis pipeline described above. For transparency and reproducibility, all analysis scripts and statistical procedures are publicly available at https://github.com/milecap/MRIMEG_pipelines.

Further details on the MRI postprocessing pipeline are given on Supporting Information 3.

## Results

### Phantom measurements: validation of SL-fMRI detection possibilities

To test the experimental feasibility of detecting a functional SL-fMRI contrast tuned to our experimental paradigm, we started with a phantom experiment tuned to create an oscillating magnetic field at 16 Hz (activation). Figure 2c shows the result of each postprocessing step, clearly highlighting the peak at the block stimulation frequency after rectification and the higher variance when dividing in on and off blocks.

### In-vivo measurements of neuronal activity with MEG

MEG data was successfully analysed for 12 out of 13 participants. The remaining participant was excluded due to incorrect acquisition timing. Figure 3a presents the results of the average frequency spectra across all subjects. MEG recordings confirmed robust stimulus-locked activity in occipital sensors during visual flicker stimulation. The time–frequency maps (Figure 3a–c) showed clear spectral peaks at 8, 16, and 24 Hz during the 6 s epochs averaged across the stimulation periods for both stimulation sides, absent during rest. The power spectra ratio between stimulation/no-stimulation periods, averaged across occipital sensors (Figure 3d), showed maximal entrainment at 16 Hz, consistent with the second harmonic of the flickering checkerboard, and a smaller peak at 8 Hz, but no response at the control frequency of 36 Hz (green arrow). Violin plots of field ratios (Figure 3e) confirmed that responses at 8 Hz and 16 Hz were consistently above zero across participants, whereas no significant response was detected at the off-resonance control frequency (36 Hz), validating this as an appropriate control frequency.

**Figure 3:**
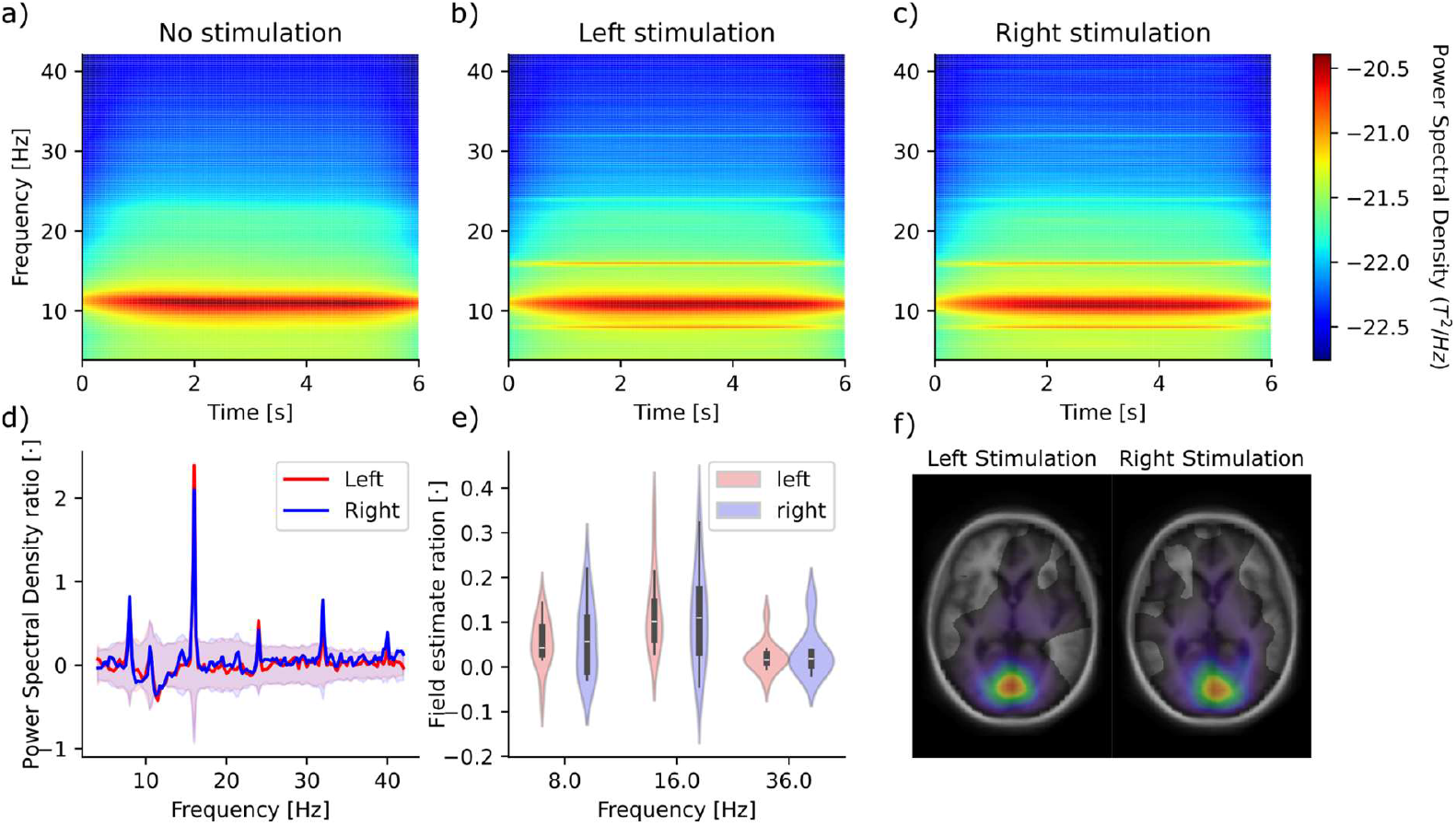
Group MEG visual stimulation results. Time–frequency maps of power spectral density from occipital sensors during a) no flashing checkerboard stimulation, b) right-, and c) left-field stimulation (Fig. 1). Clear spectral peaks appear at 8, 16, and 24 Hz during stimulation, absent in the rest condition (logarithmic color scale). d) Ratio (stimulation vs rest) of power spectral densities of occipital sensors, showing power peaks at 8, 16, 24 and 36 Hz (stimulation frequencies and harmonics). e) Violin plots of field ratios (stimulation – no stimulation, mean-centered). Ratios are significant at 8 Hz and 16 Hz, but not at 36 Hz (control), consistent with stimulus-locked activity. f) Statistically thresholded DICS activation maps averaged for all subjects for the ratio between stimulation and no stimulation in the left and right conditions.

### In-vivo measurements of neural currents with SL-fMRI

Figure 4 illustrates representative normalized activation maps at the block-stimulation frequency (0.0625) before and after RFR processing. Prior to RFR, all sequences show apparent activity in the visual cortex at the stimulation frequency, reflecting BOLD effects. Subject 1 (right-field stimulation) and Subject 2 (left-field stimulation) show correct lateralization of the pre-RFR activity. After RFR, most activations disappear, with some contrast remaining in the Bal-REX on-resonance (16 Hz) map of Subject 1, while no activation persists in the Bal-SIRS, off-resonance controls, or BOLD maps. In Subject 2, all activations are fully suppressed after RFR. This shows that the RFR procedure can correctly eliminate the hemodynamic effect. Nevertheless, contrary to the phantom experiments tuned at the stimulation frequency of 16 Hz, and despite the confirmation of this frequency activation from the MEG results, the in-vivo results show no consistent SL-fMRI induced contrast.

**Figure 4:**
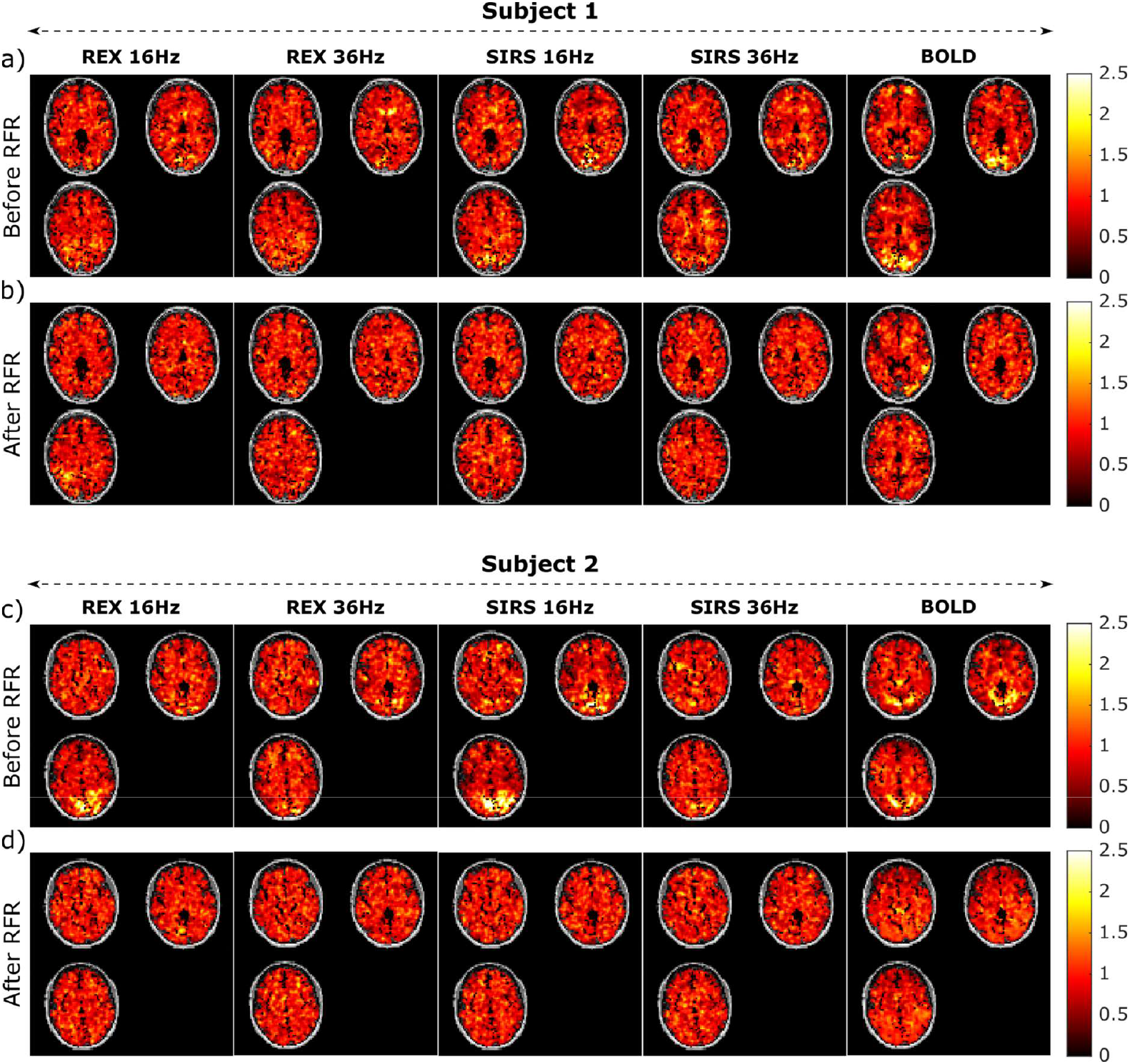
Regression-Filtering-Rectification effects on visual activation maps from Spin-Lock fMRI. Unthresholded flashing checkerboard activation maps at the stimulation frequency (0.0625 Hz) are shown before (a, c) and after (b, d) the regression–filtering–rectification (RFR) procedure for on-resonance Bal-REX (16 Hz), off-resonance Bal-REX (36 Hz), on-resonance Bal-SIRS (16 Hz), off-resonance Bal-SIRS (36 Hz), and conventional BOLD EPI in two representative subjects. Maps were normalized in frequency and space aligned to a common scale for visualization.

Despite scattered highlighted areas after RFR (as for example Subject 1 Figure 4), neither Bal-REX nor Bal-SIRS showed consistent activation at 16 Hz relative to control frequencies. Power spectra within anatomically defined V1 ROIs (Figure 5) lacked peaks at the stimulation frequency following RFR processing, both at individual and group levels.

**Figure 5:**
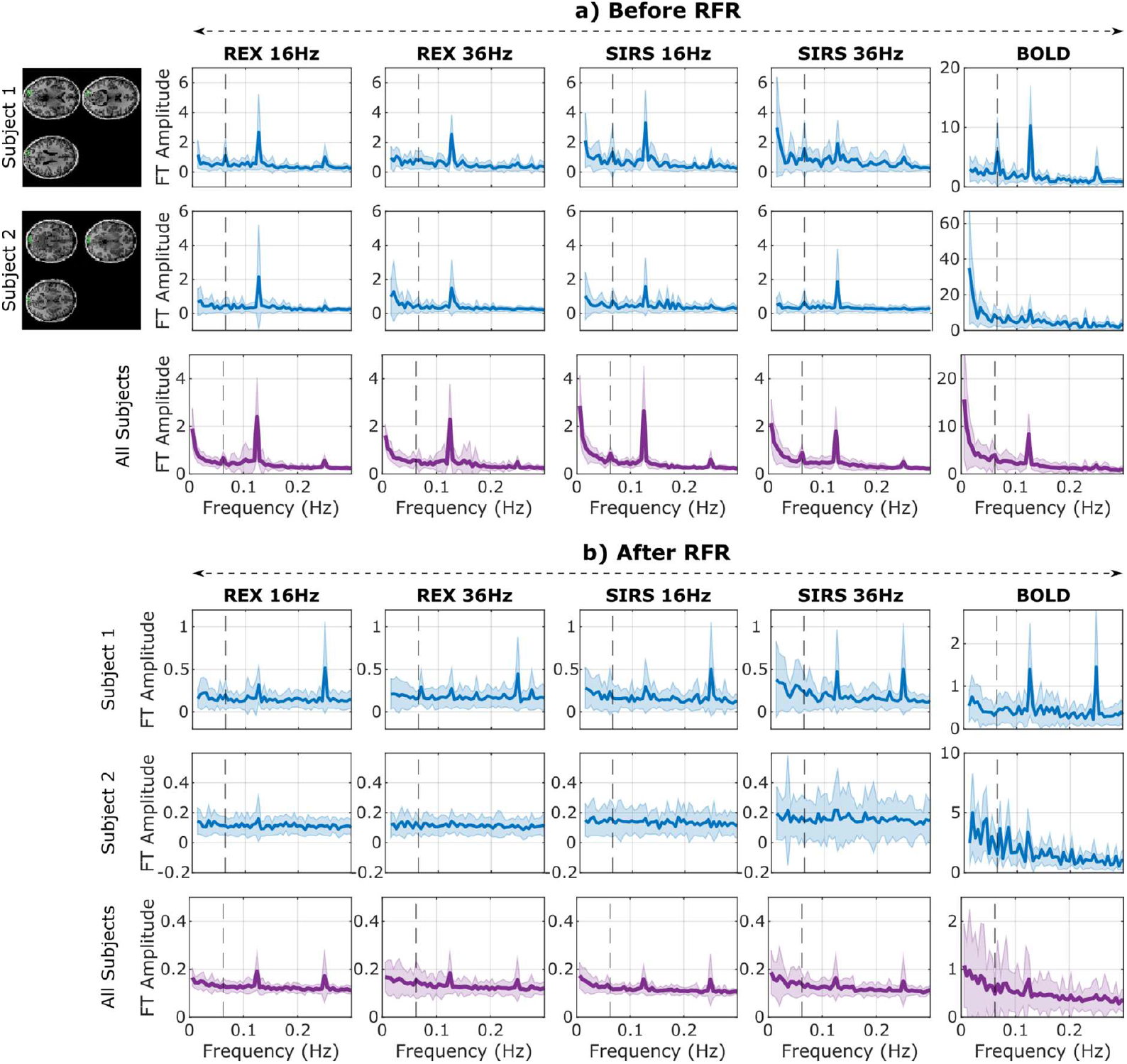
Spin-Lock fMRI power spectra in the visual cortex. Frequency-domain data from two representative subjects and the group average within the anatomically defined visual cortex, shown (a) before and (b) after the regression–filtering–rectification (RFR) procedure. Results are presented for on-resonance Bal-REX (16 Hz), off-resonance Bal-REX (36 Hz), on-resonance Bal-SIRS (16 Hz), off-resonance Bal-SIRS (36 Hz), and conventional BOLD EPI acquisitions. The vertical dashed line at 0.0625 Hz marks the expected block-stimulation frequency target.

Because spin-lock contrast is expressed in terms of signal variance, Figure 6 reports both the individual and group-average distribution of signal variability across the 13 participants. Contrast was computed as the standard deviation of the functional signal within the primary visual cortex (V1), defined according to the Destrieux atlas for the individual subjects for a) Ba-REX, b) Bal-SIRS and c) averaged across subjects for side 1. No significant differences in variance were observed between stimulation and rest periods for either the spin-lock or BOLD acquisitions. In addition, the standard deviation of the BOLD signal was consistently higher than that of the spin-lock signal. No BOLD activation survived the applied preprocessing and statistical thresholds.

**Figure 6:**
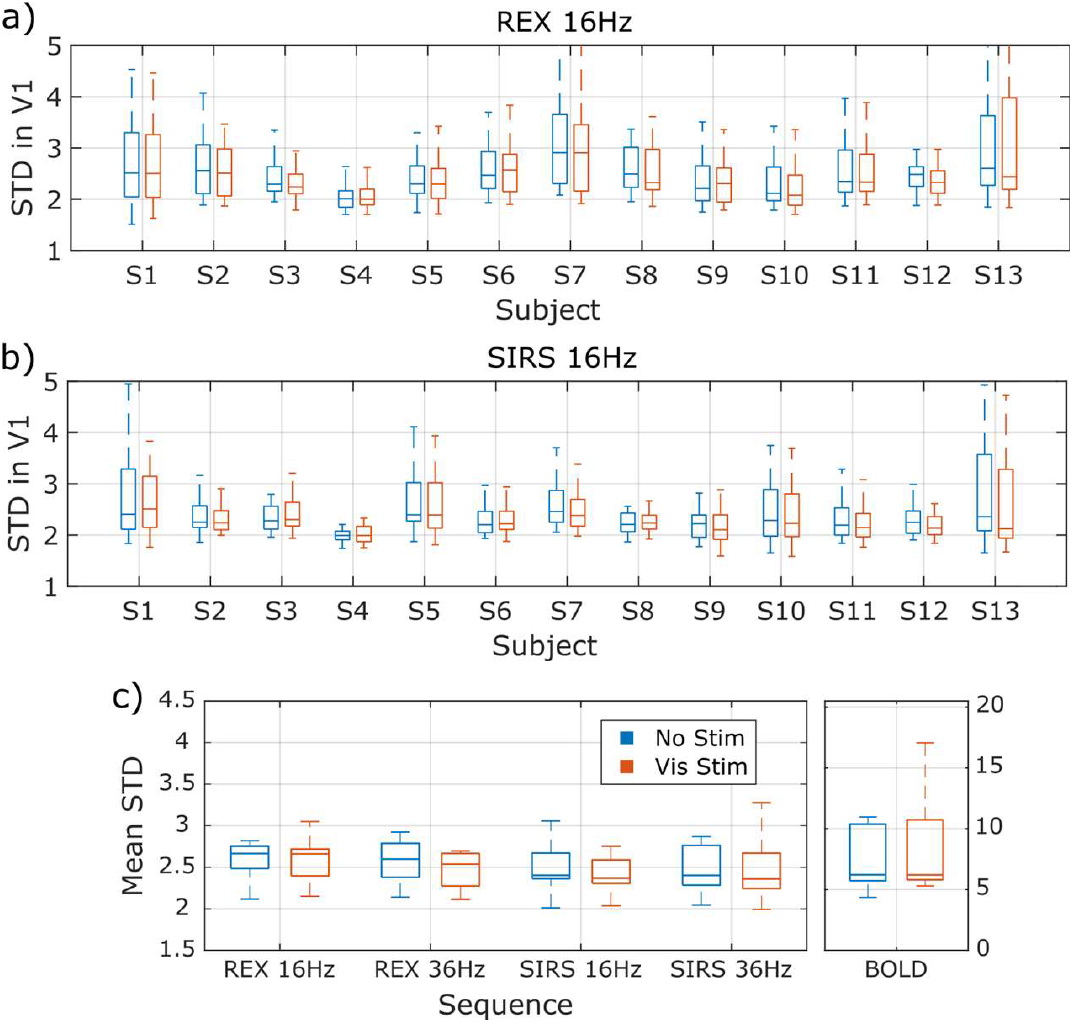
Group results of spin-lock fMRI in visual cortex. Mean standard deviation for the left-side-stimulated and non-stimulated periods within the visual cortex for a) Bal-REX, b) Bal-SIRS, and c) averaged across all subjects and for each sequence: on-resonance Bal-REX (16 Hz), off-resonance Bal-REX (36 Hz), on-resonance Bal-SIRS (16 Hz), off-resonance Bal-SIRS (36 Hz), and conventional BOLD EPI acquisitions.

**Figure 7:**
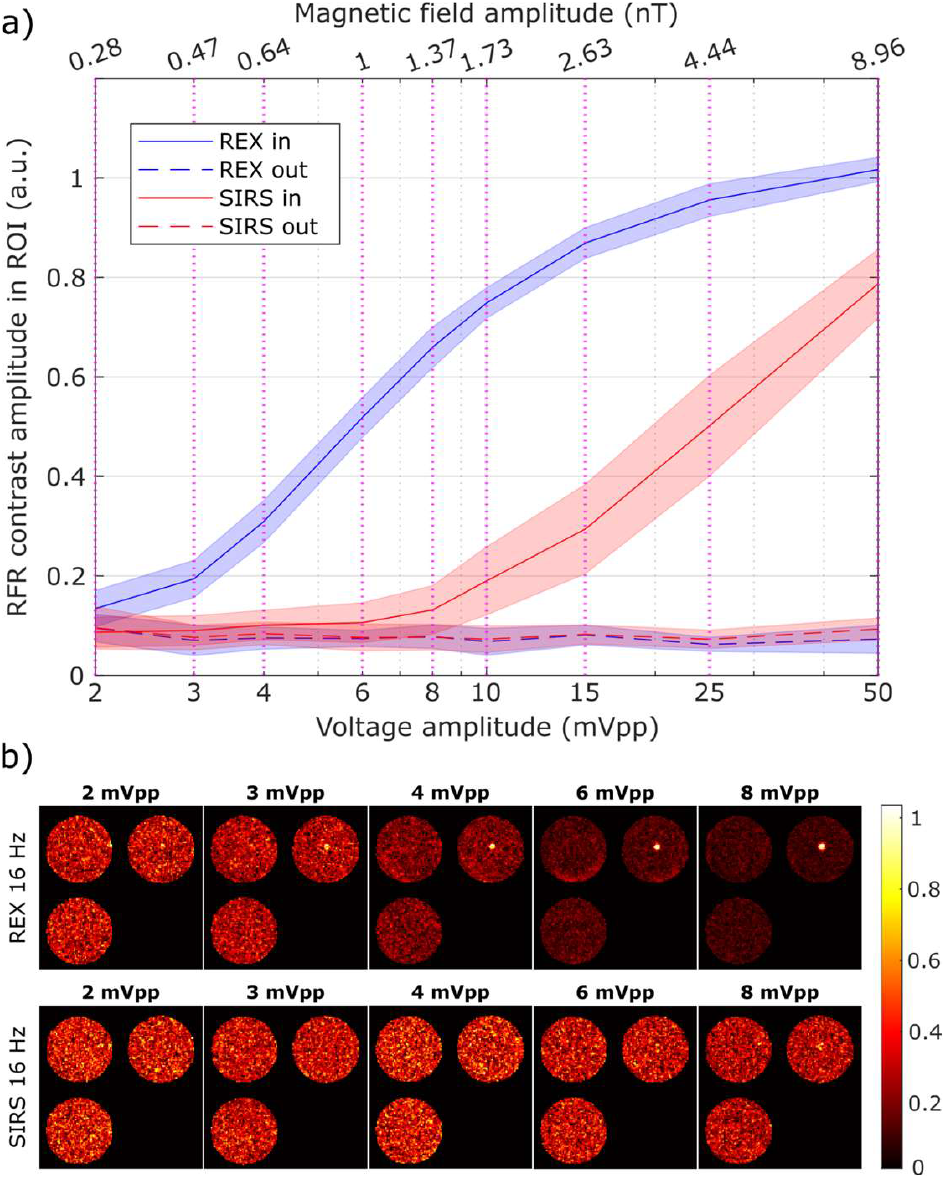
Phantom estimation of Spin-Lock fMRI sensitivity limits. a) RFR contrast as a function of the voltage amplitude for a ROI inside (in) and outside the phantom loop coil (out) for both SIRS and REX acquisitions. A t-test comparing the contrast distribution between the in and out ROIs revealed significant differences (p<0.01) in REX until 0.28 nT and SIRS until 0.6 nT. b) RFR contrast maps for REX and SIRS signals as a function of the applied field’s amplitude show both sequences’ localization capabilities.

We observed no systematic differences across alternative analysis configurations or stimulation side (see Methods), and none of the tested parameter variations yielded consistent spin-lock–specific activation.

### Sensitivity limits of spin-lock fMRI to detect oscillating currents

Having established basic feasibility with our first phantom tests (Figure 2) and verified the dominant neural oscillation frequency via MEG recordings (Figure 3), we were unable to reproduce consistent spin-lock–based contrast in vivo (Figures 4 to 6). To assess whether this failure reflects physiological limits or methodological sensitivity constraints, we performed further phantom experiments to estimate the minimum detectable oscillatory current amplitude with spin-lock fMRI and to compare this threshold to the average biomagnetic field strengths observed in MEG.

Using the MEG-based estimation procedure, we quantified the stimulus-locked magnetic field amplitudes in occipital cortex during visual stimulation. At the stimulation frequency of 16 Hz, left- and right-field stimulation elicited an estimated group-average local field amplitude of 0.073 ± 0.006 nT. To complement this measurement, we systematically reduced the amplitude of a 16 Hz alternating current applied to the phantom coil and evaluated the resulting spin-lock fMRI signal for both sequences. These experiments confirmed the higher functional sensitivity of the Bal-REX acquisition relative to Bal-SIRS, particularly as the imposed magnetic field approached the detection limit. We estimated sensitivity thresholds of approximately 0.2 nT for the Bal-REX sequence and 0.6 nT for the Bal-SIRS sequence. Results shown correspond to the RFR analysis pipeline, with comparable findings obtained using the SVarM pipeline.

Together, these findings indicate that the measured detection limits of the spin-lock acquisitions in our experimental configuration were ∼3-fold (REX) to 9-fold (SIRS) higher than the average fluctuating field amplitude estimated from MEG. This sensitivity gap likely contributed to the lack of in-vivo spin-lock fMRI effects observed here.

## Discussions

This study systematically evaluated the in-vivo sensitivity of spin-lock fMRI to detect neuronal magnetic fields in humans, using BOLD-fMRI and MEG as benchmarks and calibrated phantom experiments on a 3T clinical MRI scanner. The main finding is that no significant SL-fMRI activation was observed during a visual flashing-checkerboard stimulation, despite robust biomagnetic and hemodynamic responses detected by MEG and BOLD. Quantitative comparison with phantom-derived detection thresholds indicates that the neuronal magnetic field amplitudes estimated by MEG (∼0.07 nT) lie below the minimal detectable levels of the implemented spin-lock sequences (≥ 0.2 nT for balanced REX and ≥ 0.6 nT for balanced SIRS) under the present acquisition conditions.

The phantom experiments confirmed that both Bal-REX and Bal-SIRS sequences operated as expected (Bal-REX with higher sensitivity than Bal-SIRS) and that the analysis procedures were capable of detecting sub-nanotesla oscillatory fields. The RFR and SVarM analysis pipelines to detect SL-fMRI functional contrast provided consistent sensitivity estimates and effectively suppressed hemodynamic and low-frequency noise in in-vivo data, supporting the methodological validity of the negative finding. While balanced REX occasionally exhibited small residual fluctuations, neither sequence produced reproducible stimulus-locked activation, preventing conclusions regarding relative in-vivo performance differences between SIRS and REX.

The MEG experiments confirmed stimulus-locked oscillatory responses at 8 and 16 Hz localized to the occipital cortex, consistent with previous visual entrainment studies. The estimated field strengths were approximately threefold lower than the minimal detectable fields in the phantom. Thus, the lack of SL-fMRI activation is quantitatively consistent with the expected physical limits of the technique.

BOLD-fMRI results revealed the expected contralateral visual activations at the expected frequency before the RFR procedure, providing further confirmation that the task design and subject compliance were appropriate and that cortical stimulation elicited genuine neuronal responses. The absence of spin-lock effects in the same regions therefore reflects an intrinsic sensitivity limitation rather than lack of stimulation or analysis failure.

A limitation of this study is the use of echo planar imaging (EPI) readout instead of the previously used spiral acquisition [Truong et al., 2019]. That study employed spiral readout and shorter (4 s) stimulation blocks, potentially shifting the dominant frequency content away from low-frequency physiological confounds and improving signal-to-noise characteristics. Although SL contrast is encoded in signal variance rather than mean intensity, spiral acquisition may provide higher effective SNR and reduced hemodynamic sensitivity compared with EPI. Systematic cross-validation of spiral and EPI readouts under identical conditions will be necessary to determine whether acquisition strategy alone can bridge the observed sensitivity gap.

Several additional factors may limit Spin Lock fMRI sensitivity in the present configuration. First, neuronal magnetic fields are vectorial and only the component parallel to B_0_ can contribute to spin-lock interaction, field orientations predominantly tangential to the cortical surface may therefore produce negligible SL effects. Second, the experiment was performed at 3T, where T_1_ρ relaxation and B_1_ inhomogeneities constrain achievable spin-lock amplitudes. Ultra-low field implementations may offer improved sensitivity by diminishing B_1_ effects. Third, the simple visual stimulation paradigm may have led to habituation or reduced attentional engagement, lowering response amplitude. Future studies using parametric modulation of stimulation frequency or intensity may help delineate detection thresholds more precisely. Finally, physiological and pathological neuronal events differ substantially in amplitude and synchrony. Although the present implementation does not achieve sufficient sensitivity to detect physiological oscillatory fields, stronger pathological signals, such as epileptiform activity, may fall within detectable ranges, suggesting potential translational applications in disorders characterized by abnormally large or hypersynchronous neuronal currents.

Future work will focus on extending the present framework to alternative readout schemes and systematic quantification of the magnetic field amplitudes and frequencies that can be detected under realistic in-vivo conditions.

## Conclusions

This study provides a multimodal evaluation of the detection limits of MR-based neural current imaging mapping using spin-lock fMRI. Despite the use of robust visual stimulation with careful control of acquisition and analysis procedures, SL-fMRI did not detect stimulus-locked neuronal activity at field strengths estimated by MEG, indicating that the current sensitivity threshold lies above the physiologically generated fields. These findings define a quantitative boundary for future methodological developments and contribute a reference for the continued search for direct MRI detection of neuronal currents.

