## Supporting information for "Exploring the sensitivity limits of neuronal current imaging with MRI and MEG in the human brain"

### Supporting information 1: Detailed MEG acquisition and analysis pipeline

#### MEG Data acquisition

We recorded MEG data in a two-layer magnetically shielded room (AK3B, Vacuum Schmelze, Hanau, Germany) using a Neuromag VectorView system (306 channels: 204 planar gradiometers, 102 magnetometers). Signals were sampled at 1 kHz with a 330 Hz anti-aliasing low-pass filter and a 0.03. Hz high-pass drift filter. We digitized head shape and anatomical landmarks (nasion, left and right preauricular points) using an electromagnetic tracking system (Polhemus Inc., Vermont, USA) and placed five head position indicator (HPI) coils to continuously monitor head position during acquisition. Moreover, at least 200 additional points have been recorded on the head surface of head subject, for later re-alignment of the MRI derived surfaces. We presented stimuli and triggers with a ProPixx system (VPixx Technologies, Dorval, Canada), ensuring 1 ms temporal synchronization between MEG triggers and the actual stimuli presentation. Eye movements were recorded monocularly at 1 kHz using an MEG-compatible EyeLink system (SR Research Ltd., Ottawa, Canada).

Each trial consisted of an 8 s flickering checkerboard stimulus (8 Hz; black/white; dark grey background, RGB = 0.3, 0.3, 0.3; subtending 17° of visual angle), followed by an 8 s rest period. Before each stimulation block, we changed the color of the fixation cross to prompt participants to blink and then maintain fixation during stimulation. We repeated the stimulation/rest cycle 36 times, presenting the checkerboard in the lower left visual field for the first 18 repetitions and in the lower right for the last 18 (Figure 1a of the main text). The full experiment was repeated twice, therefore there were 36 left and right stimulation periods and 64 resting periods. Additionally, before the recording session, at least 5 minutes of empty room recording (sensors only) have been recorded.

#### MEG Preprocessing:

We performed all MEG analyses using MNE-Python (Gramfort et al., 2014) and custom Python 3.10 scripts.

We visually inspected raw MEG data to identify and exclude low-quality channels and segments contaminated by SQUID noise, muscle activity, or other artifacts. Empty-room recordings were similarly inspected, and bad channels and segments were manually marked. We processed data using the MEGIN MaxFilter implementation of Signal Source Separation (Taulu & Simola, 2006) with temporal extension (correlation threshold = 0.98; sliding window = 10 s) to suppress environmental noise and correct for head movements.

Eye-tracking enabled automatic detection of ocular artifacts, followed by manual verification. Segments containing blinks, saccades, or missing fixation were discarded. Residual noise was removed using Independent Component Analysis (Extended Infomax ICA; Lee et al., 1999), with visually identified artifactual components removed. On average, 25+- 9 segments were retained for left stimulation, 19 +- 11 for right stimulation, and 38 +- 18 for rest, with a minimum of 5 epochs (corresponding to 40 seconds of data) available across subjects and conditions.

#### MEG Analysis:

We computed power spectra separately for magnetometers and planar gradiometers in each subject and condition using a multitaper method with Discrete Prolate Spheroidal Sequences. Spectra were estimated from 4 to 42 Hz with 0.25 Hz resolution, using a 2 Hz bandwidth to average three tapers per frequency. For the time frequency estimation, spectra were computed, at each 20 ms time window, with the same parameters. Sensor spectra were then pooled together and averaged, per each condition, across subjects. Assuming the activity is mostly generated in the occipital region, a subset of occipital sensors was selected for visualization. Resulting spectral power density and spectral time-frequency maps are shown in Figure 3a-d.

In order to validate the occipital source of activity, DICS activation maps have been computed for each subject and experimental condition. For each subject, we defined both a volumetric source model (grid spacing: 3mm) and head/skull/brain enclosing mesh surfaces starting from individual T1w MRI scans (REF FreeSurfer). Surfaced and volumetric source model have been co-registered to the sensors coordinate frame by manually aligning the landmarks (left/right pre auricolar points and nasion) to the T1w data, and then refining automatically the alignment using the additional digitised points. For each subject, we checked manually the quality of the alignment and of the surface reconstructions. This led to the exclusion of a participant from further source level analyses, since the corresponding skull and head surfaces were not correctly extracted. Starting from the surfaces, we computed a BEM forward model and, then, a set of DICS beamforming inversion filters. For DICS filters computation, we at first computed the cross spectral density (CSD) pooling all the conditions together: this CSD has been used as signal covariance. Moreover, as noise covariance for the DICS algorithm, we used the CSD from the emptyroom recordings, preprocessed with the same SSS spatial filter as the actual data. After obtaining the common DICS filters, we computed activations by applying them to condition specific cross spectral density matrices, thus obtaining, for each subject, a left/right stimulation and a rest condition volumetric activation map. Finally, we morphed each individual map to a common space (the FreeSurfer average) for further statistical and group analysis. From the common space morphed maps, we computed the ratio between the activation in the left/right stimulation condition and the rest condition. These ratios were tested against 1 using a permutation approach. We ran a group level statistical bootstrap test (5000 permutations; T-test statistics), finally obtaining two maps of activation ratio (left/right stimulation vs rest). The activation maps, for the most prominent frequency of 16 Hz, are shown in Figure 3f.

Finally, for each subject and for each condition, an estimate of the local magnetic field strength have been obtained following the approximation in Bodurka et al. (2002). We filtered the magnetic field time series as obtained by occipital magnetometers only, in a narrow band around each of the following frequencies: 8 Hz, 16 Hz and 36 Hz (the latter being a control frequency where no effect of the periodic stimulation is expected). Then the Hilbert transform was computed and the local field estimate obtained by multiplying the average Hilber envelope for a factor $10^3$. This led to an average value of the local field at the strongest peak (16 Hz) of 0.073 ± 0.006 nT. Finally, the ratio of the local field between the left/right stimulation condition and the rest was computed for each subject and shown, as a violin plot distribution, in Figure 3e.

#### Supporting information 2: Bloch Simulations and Optimization of Spin-Lock Duration.

To determine the optimal spin-lock duration (TSL) for in-vivo experiments, we relied on numerical simulations of the Bloch equations previously described in Capiglioni et al. (2022) and Capiglioni et al. (2023). These simulations modelled the dependence of spin-lock–induced signal modulation on both TSL and the initial phase of an oscillatory target magnetic field under resonance conditions.

Simulations were performed under on-resonance conditions (FSL = 16 Hz) and assumed sinusoidal target fields with fixed frequency of 16 Hz and varying amplitudes in the short nT range. To account for the unknown initial phase of physiological oscillations, contrast estimates were averaged across all possible initial phases with 100 points simulated uniformly between 0 and 2 pi.

Figure S1 illustrates the simulated standard deviation of the spin-lock signal as a function of TSL for balanced SIRS (panel a) and balanced REX (panel b) at a target frequency of 16 Hz. Curves are shown for increasing target field amplitudes (BNC = 0–0.05 nT). The dashed vertical line indicates TSL = 112 ms. For both sequences, signal modulation exhibits a pronounced dependence on TSL, with maximal variance occurring near 100–130 ms.

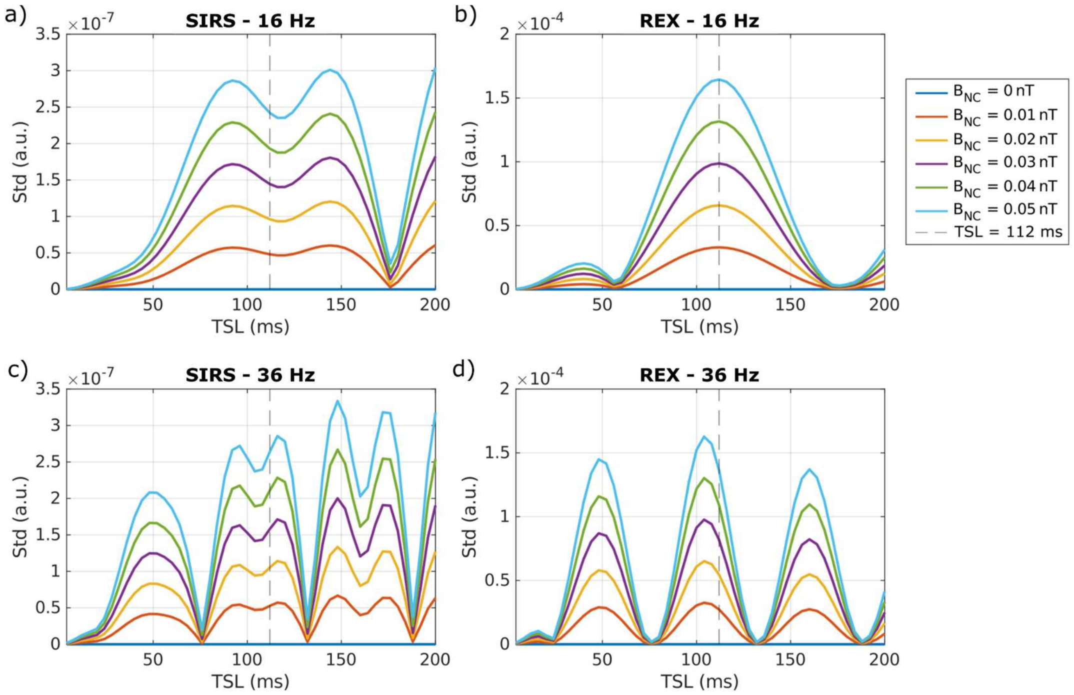


***Figure S1: Simulated dependence of spin-lock signal modulation on spin-lock duration (TSL)****. Standard deviation of the simulated signal as a function of TSL at 16 Hz for balanced SIRS (a) and balanced REX (b) and at 36 Hz for SIRS (c) and REX (d) . Curves correspond to increasing amplitudes of the oscillatory target field (BNC = 0–0.05 nT). The dashed vertical line marks the selected TSL of 112 ms.*

To corroborate that the observed contrast is due to the rotary saturation effect, measurements with off resonance SL preparation were also conducted. Given that stimulation at a specific frequency has been shown to elicit activations at harmonics (in this case 16, 32, and 40 Hz), the off-resonance control frequency was set to 36 Hz. The contrast was therefore also optimized at the off-resonance frequency. Figure S1 displays the response of the std for SIRS (panel c) and REX (panel d) on resonance at 36 Hz

(FSL = 36 Hz with sinusoidal field at 36 Hz). Across simulated field amplitudes and frequencies, TSL = 112 ms consistently provided near-maximal signal variation while preserving stable $T_1\rho$ weighting and frequency selectivity. Based on these results, a TSL of 112 ms was selected for all in-vivo and phantom experiments to maximize sensitivity to oscillatory magnetic fields while maintaining comparable relaxation weighting across acquisition conditions.
